# Never in Mitosis Kinase 2 regulation of metabolism is required for neural differentiation

**DOI:** 10.1101/2022.08.23.504995

**Authors:** Danielle M. Spice, Tyler T. Cooper, Gilles A. Lajoie, Gregory M. Kelly

## Abstract

Wnt and Hh are known signalling pathways involved in neural differentiation and recent work has shown the cell cycle regulator, Never in Mitosis Kinase 2 (Nek2) is able to regulate both pathways. Despite its known function in pathway regulation, few studies have explored Nek2 within embryonic development. The P19 embryonal carcinoma cell model was used to investigate Nek2 and neural differentiation through CRISPR knockout and overexpression studies. Loss of Nek2 reduced cell proliferation in the undifferentiated state and during directed differentiation, while overexpression increased cell proliferation. Despite these changes in proliferation rates, Nek2 deficient cells maintained pluripotency markers after neural induction while Nek2 overexpressing cells lost these markers in the undifferentiated state. Nek2 deficient cells lost the ability to differentiate into both neurons and astrocytes, although Nek2 overexpressing cells enhanced neuron differentiation at the expense of astrocytes. Hh and Wnt signaling were explored, however there was no clear connection between Nek2 and these pathways causing the observed changes to differentiation phenotypes. Mass spectrometry was also used during wildtype and Nek2 knockout cell differentiation and we identified reduced electron transport chain components in the knockout population. Immunoblotting confirmed the loss of these components and additional studies showed cells lacking Nek2 were exclusively glycolytic. Interestingly, hypoxia inducible factor 1α was stabilized in these Nek2 knockout cells despite culturing them under normoxic conditions. Since neural differentiation requires a metabolic switch from glycolysis to oxidative phosphorylation, we propose a mechanism where Nek2 prevents HIF1α stabilization, thereby allowing cells to use oxidative phosphorylation to facilitate neuron and astrocyte differentiation.

## 2. Introduction

Hedgehog (Hh) and Wnt signalling pathways are vital for neural development and are involved during *in vivo* and *in vitro* neural differentiation [1–5]. During neurulation the notochord and floor plate secrete Hh, inducing ventral neural phenotypes, and the roof plate secretes Wnt, inducing dorsal neural phenotypes [6–8]. These signals not only pattern the dorsal-ventral axis of the neural tube but are also vital for patterning its anterior-posterior axis [7] and defining the midbrain-hindbrain boundary [9,10]. Activation of the Hh pathway occurs when the Hh ligand binds to its receptor Patched (Ptch), which relieves the inhibition of Smoothened, allowing Smoothened to translocate to the plasma membrane [11]. Translocation causes the Suppressor of Fused (SUFU) and Gli complex to be recruited to the primary cilium where SUFU dissociates from the full-length form of Gli, which enters the nucleus to activate transcription [12,13]. In the absence of ligand, SUFU-bound Gli is either sequestered in the cytoplasm or is targeted for partial proteosomal degradation converting Gli to a transcriptional repressor [8,14]. In the case of the Wnt pathway, it is either canonical and involves β-catenin or non-canonical and linked to planar cell polarity or Ca^2+^ signalling. In either case, one of 19 vertebrate Wnt ligands binds the transmembrane receptor Frizzled (Fzd), which signals to Dishevelled (DVL) and in the canonical pathway causes the recruitment of the destruction complex [15–17] that includes Axin, adenomatous polyposis coli, glycogen synthase kinase 3 (GSK3), casein kinase 1, and β-TrCP, a ubiquitin ligase [15,18]. This recruitment and the destabilization of Axin attenuates the hyper-phosphorylation of β-catenin by GSK3 allowing β-catenin cytoplasmic accumulation and translocation to the nucleus where it interacts with TCF/LEF transcription factors inducing Wnt target gene transcription [19].

Recently, Never in Mitosis Kinase 2 (Nek2) was reported as a novel regulator of both canonical Wnt and Hh pathways [20–23], but few studies have explored its role in embryonic development. What is known, however, is that Nek2 plays an essential role in left-right asymmetry of the Xenopus heart [24], its overexpression in the Drosophila eye ectopically induces *wingless* (*Wnt1*) signalling and reduces retinal neuron differentiation [25], and in mouse, Nek2 expression is reported in the brain of the E10.5 embryo [26]. The role of this protein in signalling, however, originated from reports of its involvement as a centrosomal linker protein during the cell cycle, showing Nek2 binding and phosphorylating DVL [20] and β-catenin [21] at centrosomes. Nek2, serving as a Gli1/2 target gene [22], also phosphorylates and stabilizes SUFU [23], placing it as negative regulator of the Hh pathway. Overexpression of Nek2 has also been linked to multiple Wnt- and Hh-associated cancers, including those causing brain malignancies [27–32].

Given that Hh and Wnt signalling are essential in neural differentiation, and that Nek2 acts as a regulator of both pathways, we sought to explore the role of Nek2 in neural differentiation using P19 embryonal carcinoma cells. These cells can be differentiated towards neurons and astrocytes in the presence of retinoic acid (RA) [33–37], which affects Wnt [3,5,38,39] and Hh [37,40] signalling. Herein, we report the presence of Nek2 throughout RA-induced differentiation and through overexpression studies and CRISPR-Cas9 ablation, demonstrate an essential role of Nek2 in neuron and astrocyte differentiation. Furthermore, we show that Nek2 overexpression is linked to increased Wnt signalling, however, proteomic analysis suggests an essential role of Nek2 outside of signalling and connects its involvement to metabolic regulation.

## 3. Methods

### 3.1. Cell Culture

Mouse P19 embryonal carcinoma cells were a generous gift from Dr. Lisa Hoffman, Lawson Health Research Institute, UWO. Cells were maintained in DMEM containing 5% fetal bovine serum (FBS, ThermoFisher) and 1% penicillin/streptomycin (ThermoFisher) on adherent tissue culture plates (Sarstedt). Wildtype and mutant cells were passaged every 4 days or when they reached a confluency of 70%, whichever occurred first. Following previously published protocols [37,41], differentiation was induced in wildtype and mutant cells by plating 1.05 × 10^6^ cells on bacterial grade Petri dishes (Falcon) in the presence of 0.5 μM retinoic acid (RA) for 4 days to form embryoid bodies (EB). These aggregates were collected and re-plated on adherent tissue culture dishes in the presence of 0.5 μM RA, which in wildtype cells, develop into neurons after 10 days and form astrocytes by day 17. For BIO-treated experiments, cells were induced to differentiate in the presence of 0.5 μM RA on days 0 and 4, 0.5 μM RA treatments on days 0 and 4 plus 10 nM BIO (6-Bromoindirubin-3’-oxime, Sigma) treatment on day 0, or 10 nM BIO treatment on day zero plus 0.5 μM RA on day 4. Cells were also treated with 10 nM BIO alone with no subsequent RA. For XAV treatments, cells were induced to differentiate in the presence of 0.5 μM RA on days 0 and 4, plus 5 μM XAV on day 0. For metabolic experiments cells were seeded and allowed to adhere for 16 hours, then treated with either media alone, DMSO, 2.5 μM Oligomycin A (BioShop) or 50 mM 2-Deoxy-D-glucose (2-DG, Sigma) for 24 hours. Cell counts and viability were measured by trypan blue exclusion. Cells for all experiments were maintained at 37°C and 5% CO_2_.

### 3.2. Preparing CRISPR plasmids

*Nek2* sgRNAs (Supplementary Table 1) were cloned into the CRISPR-Cas9 plasmid pSpCas9(BB)-2A-Puro (PX459) V2.0 (Addgene plasmid 62988; http://n2t/addgene:62988) using the protocol by Ran et al. 2013 [42]. Prior to cloning, sgRNAs were amplified and phosphorylated, then subsequently digested with the BbsI restriction enzyme along with the vector. Digested sgRNA and vector were incubated at room temperature for 1.5 hours in the presence of T7 ligase (New England Biolabs). Plasmids were transformed into transformation competent DH5alpha *E. coli* cells and positive colonies were collected using a Mini Prep kit (Qiagen) and sequenced using the U6 primer (Supplementary Table 1) at the London Regional Genomics Centre (Robarts Research Institute, London, ON).

### 3.3. Generating Nek2 deficient cell lines

Two μg of sgRNA containing plasmid was incubated with 10 μL of Lipofectamine 2000 (Invitrogen) for 20 minutes and then added to high confluency P19 cells for 4 hours. Culture media was replaced, and cells were allowed to reach 70% confluency. Cells were then passaged to a 1 cell/50 μL concentration into a 96 well plate where they were incubated daily in 1 μg/mL puromycin (BioShop) selection medium for 4 days. After selection, cells were allowed to recover in compete medium until reaching 70% confluence. Mutant genotypes were identified by collecting genomic DNA (Qiagen DNeasy Blood & Tissue kit, 69504), which was amplified using PCR (DreamTaq Master Mix (2X), Thermo Scientific, K1081), a vapo.protect Thermocycler (Eppendorf) and site-specific primers (Supplementary Table 1). Amplified DNA was sequenced at the London Regional Genomics Centre and pair-wise alignments were performed using Geneious 2021 ™. Additional analysis of in/dels was performed using Synthego Performance Analysis, ICE Analysis. 2019. v3.0. Synthego; [Jan. 13, 2022].

### 3.4. Generating Nek2 Overexpression Cell Line

Two μg of plasmid was incubated in the presence of 10 μL of Lipofectamine 2000 for 4 hours. Media was changed and cells were allowed to grow to 70% confluency before being passaged. Cells were incubated in selection media containing either 1 mg/mL hygromycin or 500 μg/mL neomycin for 2 weeks, with selection media being changed every 48 hours, to generate stable lines. Plasmids used include: 1436 pcDNA3 Flag HA (Addgene plasmid #10792) and pCMV3-human NEK2-Myc (HG10054-CM, Sino Biological).

### 3.5. RT-qPCR

Real time reverse transcriptase PCR analysis was used to determine relative expression levels of various genes accompanying differentiation in wildtype and mutant cell lines. In wildtype cells, mRNA was collected at days 0, 1, 2, 3, 4, 6, 8 and 10 after RA treatment and in wildtype and mutant cells, mRNA was collected at days 0, 10, 14 and 17 after RA treatment using QIAshredder (Qiagen, 79654) and RNeasy (Qiagen, 74104) kits. RNA was reverse-transcribed into cDNA using a High-Capacity cDNA Reverse Transcription Kit (ThermoFisher Scientific, 4368814) and was amplified using the following primers: *L14, Nek2, Gli1, Ptch1, Ascl1, Dab2, Dkk1, c-Myc, Nanog* and *Oct4* (Supplementary Table 2). PCR reactions containing 500 nM each of reverse and forward primers, 10 μL of SensiFAST SYBR No-ROX Mix (FroggaBio, BIO-98050) and 1 μL of cDNA, were run using a CFX Connect Real-Time PCR Detection System (Bio-Rad). Samples were analyzed using the comparative cycle threshold method (2^-ΔΔCt^) using *L14* as an expression control and comparing normalized expression values to undifferentiated wildtype cells to determine fold changes in expression.

### 3.6. Immunoblotting

Cells at various stages of differentiation in both wildtype and mutant lines were lysed using 100-500 μL of 2% sodium dodecyl sulfate buffer with 10% glycerol, 5% 2-mercaptoethanol, 1:200 of 1X Halt Protease Inhibitor Cocktail (ThermoFisher Scientific, 1862209), at pH 6.8. Proteins were sonicated on ice for 30 seconds and quantified using a DC™ Protein Assay (Bio-Rad, 5000113) before loading 10 μg of protein to 6-10% polyacrylamide gels. Proteins were separated for approximately 1.5 hours at 120V, and then transferred overnight at 20V to PVDF membranes (Bio-Rad, 1620177). Membranes were placed for 1 hour at room temperature in 5% skim milk in Tris-buffered saline containing 0.1% Tween-20 (TBS-T). After washing, blots were incubated overnight at 4°C in the following primary antibodies: β-III-Tubulin (TUJ1, 1:1000; Cell Signaling Technology, 5568), SUFU (1:1000; abcam, ab28083), GFAP (1:1000; Invitrogen, 14-9892-80), Nek2 (Santa Cruz Biotechnology, sc-55601), Dab2 (Cell Signaling Technology, 12906), ERRB (1:1000; Novus Biologicals, PP-H6705-00), Total OXPHOS (1:1000; acbam, ab110413), TOM20 (1:1000; Cell Signaling Technology, 42406), HIF1α (ab179483) and β-actin (1:10,000; Santa Cruz Biotechnology, sc-47778). Blots were washed and then incubated for 2 hours at room temperature in host-specific HRP-conjugated secondary antibody (1:10,000, Sigma) in TBS-T containing 5% skim milk. After secondary antibody incubation and washing, blots were incubated in Immobilon^®^ Classico Western HRP Substrate (Millipore, WBLUC0500) and imaged using a Chemi Doc Touch System (Bio-Rad). Densitometry analysis was performed using ImageLab Software (Bio-Rad).

### 3.7. Immunofluorescence and Phase Contrast Imaging

Wildtype and *Nek2* knockout cells were treated with 0.5 μM RA as described [37] for 17 days on poly-L-lysine hydrobromide (Sigma, P5899) coated coverslips. Cells on coverslips were washed with PBS for 1 minute and then fixed with 4% paraformaldehyde in PBS for 10 minutes at room temperature. Ice-cold PBS was used to wash coverslips, then cells were permeabilized with 0.2% Triton-X-100 in PBS for 10 minutes at room temperature. Cells were again washed before being blocked for 30 mins at room temperature in PBS containing 1% bovine serum albumin, 22.52 mg/mL glycine and 0.1% Tween-20. Coverslips were then incubated overnight in a humidity chamber with the following primary antibodies: β-III-Tubulin (TUJ1, 1:400; Cell Signaling Technology, 5568) and GFAP (1:500; Invitrogen, 14-9892-80). After primary incubation, cells were washed and incubated with secondary antibodies in blocking solution for 1 hour at room temperature. The secondary antibodies were Alexa Fluor™ 660 Goat anti-Mouse IgG (Invitrogen, A21054) and Goat anti-Rabbit IgG Alexa Fluor Plus 488 (Invitrogen, A32731TR). Finally, coverslips were washed and mounted to slides using Slowfade™ Gold Antifade Mountant with DAPI (Invitrogen, S36942) and sealed with nail polish. Images were captured on a Zeiss AxioImager Compound Fluorescence Microscope (Integrated Microscopy Facility, Biotron, Western University, London, ON).

Wildtype and *Nek2* mutant cells were treated with RA as described above, and after 4 days, EBs were imaged using a Leica DMIL LED microscope or a Zeiss inverted microscope and an OPTIKA Microscopes C-B5 camera with OPTIKA Lightview software. EB number per image area, EB area and aspect ratio were measured using ImageJ.

### 3.8. Cell Count and Viability Assays

Trypan blue exclusion assays were performed on wildtype and mutant cells in the undifferentiated state, after 4 days of RA treatment and after 2-DG or Oligomycin A treatment. In the undifferentiated state, 15,000 cells were plated on adherent tissue culture dishes. Every 24 hours for 96 hours, cells were trypsinized and a 1:1 cell suspension to trypan blue ratio was added and total number of cells counted using a CellDrop BF Brightfield Cell Counter (DeNovix). For RA treatment, cells were processed as described previously [37] and EBs were collected after 4 days. EBs were vigorously resuspended in complete media, before being counted in a 1:1 cell suspension to trypan blue ratio. Total cell number was counted, and viability was compared to the number of dead cells.

### 3.9. Caspase 3 Activity Assay

Caspase 3 activity was measured using the Caspase-3 Colorimetric Activity Assay Kit (APT131, Millipore). Briefly, cells were treated with RA for 4 days to form EBs, and then collected and lysed according to manufacturer’s instructions. Caspase-3 activity was determined through the detection of p-nitroaniline (pNA) in cytoplasmic lysates after incubation with a Caspase-3 specific substrate (Ac-DEVD-pNA). Readings were performed using a Modulus™ II Microplate Multimode Reader (Turner Biosystems) at 450nm, where background detection was subtracted and normalized to wildtype Caspase-3 activity.

### 3.10. Luciferase Assay

Wildtype and *Nek2* mutant cells were transfected with 4 μg total of DNA using 10 μL of Lipofectamine 2000. Transfected plasmids included pRL-TK (Promega) alone, or pRL-TK and pGL3-BARL (β-catenin activated reporter luciferase, Promega) or pRL-TK and pGL3-8xGli (a gift from Philip Beachy, Stanford University). Following transfection, cells were incubated for 24 hours before passaging into a 96-well plate. Cells were allowed to adhere for 48 hours before the Dual-Gli^®^ Luciferase Assay System (Promega, E2920) was used according to manufacturer’s instructions. Briefly, Dual-Glo reagent was incubated in each well containing growth media for 15 minutes before measuring Firefly luminescence (from pGL3-BARL and pGL3-8xGli plasmids). Subsequently, Dual-Glo Stop & Glo reagent was added to each well and was incubated for 15 minutes before measuring *Renilla* luminescence (from pRL-TK plasmid). To calculate initial luminescence values, background luminescence was subtracted from each well, where Firefly luminescence was divided by *Renilla* luminescence for each treatment. Relative luminescence was determined by dividing each well by the luminescence from WT cells transfected with pRL-TK only.

### 3.11. LC-Mass Spectrometry

Cells were differentiated as described, and cells were either centrifuged as EBs for collecting cells at days 1 and 4 of RA treatment or were trypsinized and centrifuged for adherent cells at days 0 and 10 of RA treatment. Lysates of P19 cells were prepared through incubation in 8 M Urea, 50 mM ammonium bicarbonate (ABC), 10 mM DTT, 2% SDS at room temperature for 10 minutes and sonicated with a probe sonicator for 20 x 0.5s pulses to shear DNA. Lysates were quantified by using a Pierce™ 660 nm Protein Assay (ThermoFisher Scientific) and stored at −80°C until future use. Lysates were reduced in 10 mM DTT for 30 min and alkylated in 100 mM iodoacetamide (IAA) for 30 min at room temperature in the dark. Next, lysate protein was precipitated in chloroform/methanol and prepared for in-solution digestion according to Kuljanin *et al.* [43]. On-pellet protein digestion was performed in 50 mM ABC (pH 8) by adding TrypLysC (Promega) at an enzyme: protein ratio at 1:25 and incubated overnight at 37°C in a ThermoMixer at 900rpm. Digestion buffer was acidified to 1% formic acid (FA) and vacuum centrifuged at 45°C. Digested peptides were desalted using in-house C18 stagetips prepared in a 200μl pipette tip. C18 stagetips were conditioned with 80% Acetonitrile (ACN)/0.1% trifluoracetic acid (TFA) followed by 5% ACN/0.1%TFA before loading and washing peptides twice with 5% ACN/0.1%TFA. Desalted peptides were eluted with 80%ACN/0.1% TFA prior to vacuum centrifugation and resuspension in 0.1% FA. Final peptide concentrations were determined by BCA quantification prior to ultraperformance liquid chromatography tandem mass spectrometry (UPLC-MS/MS).

500ng of peptides per sample was analyzed by using an M-class nanoAquity UPLC system (Waters) connected to a Q Exactive mass spectrometer (Thermo Scientific). Buffer A consisted of Water/0.1% FA and Buffer B consisted of ACN/0.1% FA. Peptides were loaded onto an ACQUITY UPLC M-Class Symmetry C18 Trap Column (5μm, 180 m x 20 mm) and trapped for 5 min at a flow rate of 10 L/min at 99% A/1% B. Subsequently, peptides were separated by were separated on an ACQUITY UPLC M-Class Peptide BEH C18 Column (130 A, 1.7 μm, 75μm × 250 mm) operating at a flow rate of 300 nL/min at 35°C using a non-linear gradient consisting of 1–7% B over 1 min, 7–23% B over 179 min, 23–35% B over 60 min, 35-98% B over 5 minutes, before increasing to 95% B and washing. Settings for data acquisition on the Q Exactive are outlined in Supplematary Table 3. Raw UPLC-MS/MS files were processed in FragPipe 1.18.0 and data analysis was performed in Perseus software (v1.16.4). Statistical significance was determined using Student’s t-test (p<0.05). Gene Set Enrichment Analysis (GSEA) using GSEA 4.2.3 was used to reference GO Biological Processes using Signal2Noise for ranking genes, all other settings were left at default.

### 3.12. Statistical Analysis

One-way ANOVA with Tukey’s post-hoc analysis was used for wildtype timeline experiments and experiments containing only 3 groups. For experiments comparing expression or protein abundance over time across two cell lines a Two-way ANOVA was used with Sidak’s multiple comparisons post-hoc analysis. All statistical analyses were performed using Prism Software (Prism version 9.3.1).

## 4. Results

### 4.1. Nek2 levels are unchanged during neural differentiation

Despite the reports showing that Nek2 regulates Hh and Wnt signalling [20–23] and both pathways are activated during P19 neural differentiation [5,37,40], few publications have explored the role of Nek2 in embryonic development [25,26,44] and to our knowledge, few limited work exists on its involvement in neurogenesis. To first address this shortfall, *Nek2* gene expression (Supplemental Figure 1) and protein levels (Fig. 1A and B) were explored throughout RA-induced differentiation, and results showed no significant change in either transcript or protein levels.

**Figure 1.**
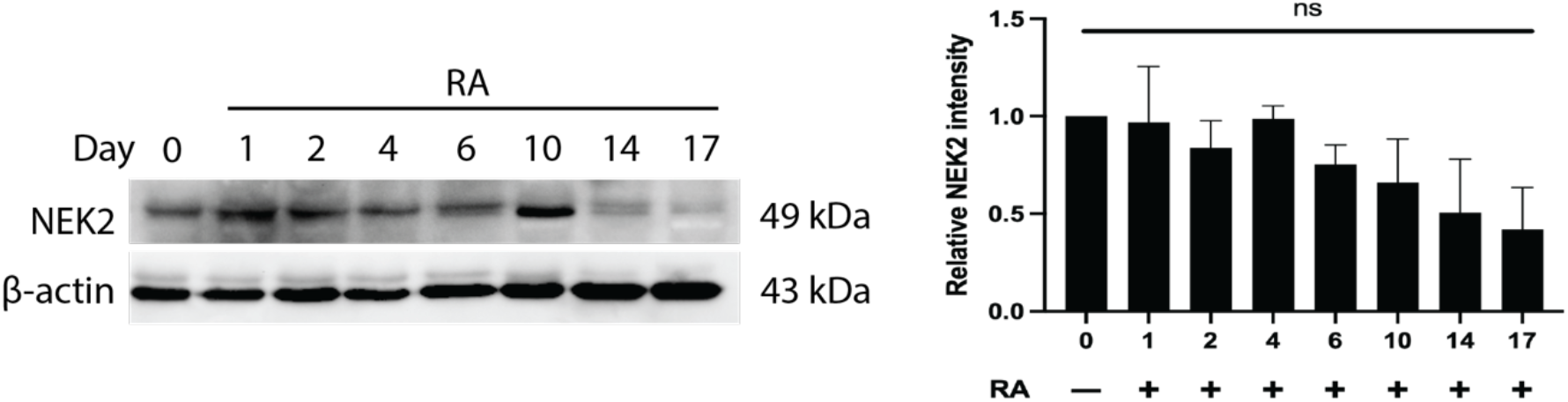
Nek2 levels do not change during neural differentiation. **(A)** Immunoblot of Nek2 at various timepoints between days 0-17 of RA treatment. **(B)** Densitometry of immunoblot in A. N=3. Bars represent mean values ±s.e.m. P-values were determined by One-way ANOVA using Tukey’s post-hoc analysis.

### 4.2. Nek2 regulates cell proliferation in undifferentiated cells

To determine the role of Nek2 in P19 cells, CRISPR-Cas9 was used to disrupt exon 2 of the coding region of the gene (gRNA sequence in Supplementary Table 1). This targeting resulted in a pool of alleles that consisted primarily of a 10 bp deletion, but also contained alleles of 1, 2, 8, 11, 17 and 18 bp deletions, as well as 5, 7 and 8 bp insertions (Supplementary Figure 2A). This polyclonal pool was labelled knockout (KO) as all alleles resulted in a frameshift mutation and immunoblotting showed no detection of Nek2 protein in these cells (Fig. 2A and B). A second gRNA, targeting exon 3 (Supplementary Table 1), resulted in a reduction in Nek2 levels (Fig. 2A and B) and identified alleles within these cells were either wildtype (WT) sequence or a 1 bp insertion (Supplementary Figure 2B). These cells were labelled as knockdown (KD) cells and although the nature of the knockdown was not explored further, it may be the result of heterozygous alleles or in defects in pre-mRNA splicing [45]. Cells were also transfected with either pcDNA or a plasmid containing human *Nek2* (pNek2-Myc), the latter showing significantly increased levels of Nek2 protein over the pcDNA control (Fig. 2A and B).

**Figure 2.**
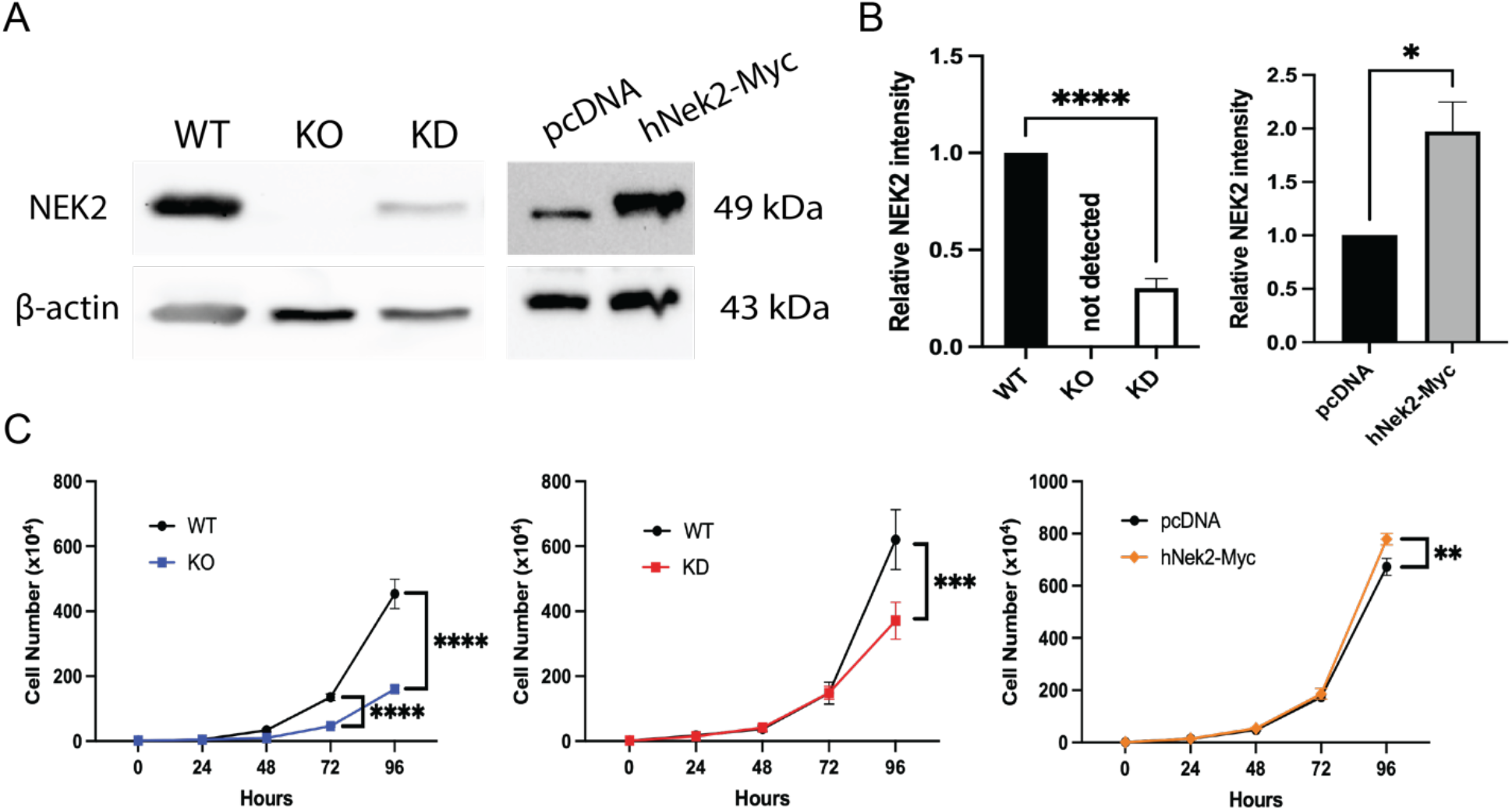
Altering Nek2 levels changes proliferation rate of undifferentiated cells. **(A)** Immunoblots of Nek2 in wildtype (WT), knockout (KO), knockdown (KD), pcDNA and hNek2-Myc transfected cells in the undifferentiated state. **(B)** Densitometry of immunoblots in A. Bars represent mean values ±s.e.m. P-values were determined by One-way ANOVA using Tukey’s post-hoc analysis. **(C)** Total cell counts of WT, KO, KD, pcDNA and hNek2-Myc transfected cells between 0-96 hours after plating in the undifferentiated state. Dots represent mean values ±s.e.m. P-values were determined by Two-way ANOVA using Sidak’s post-hoc analysis. *\*P<0.05, **P<0.01, ***P<0.001, ****P<0.0001*.

As Nek2 is a known regulator of the cell cycle [46], cell counts were performed in undifferentiated cells, and results show that the Nek2 KO caused significantly fewer cells to be present after 72 and 96 hours (Fig. 2C); KD cells had significantly fewer cells only after 96 hours (Fig. 2C). *Nek2*-overexpressing cells showed the opposite results, with more cells present than the control at 96 hours (Fig. 2C). Thus, Nek2 loss resulted in decreased cell proliferation while its overexpression caused the opposite effect.

The reduction in cell number in KO and KD cells was also observed after 4 days of RA treatment (Supplemental Figure 3A) and resulted in dramatically reduced EB size (Fig. 3A and D) and organization (Fig. 3B), without altering the number of observed EBs (Fig. 3C and D). These changes in RA treated EBs appeared to be the result of a maintained reduction in proliferation (Supplemental Figure 3A) without affecting caspase 3 activity (Supplemental Figure 3B) or viability (Supplemental Figure 3C). Significantly, *Nek2* overexpressing cells compared to the control also showed reduced EB size (Fig. 3A and D), while showing increased organization and an aspect ratio closer to 1 (Fig. 3B). Unlike the KO and KD cells, however, pNek2-Myc transfected cells had more EBs per image area compared to pcDNA transfected cells (Fig. 3C and D). Together, these data show that Nek2 is important for regulating the cell cycle in the undifferentiated state, and it appears to continue this regulation during the first days of neural induction.

**Figure 3.**
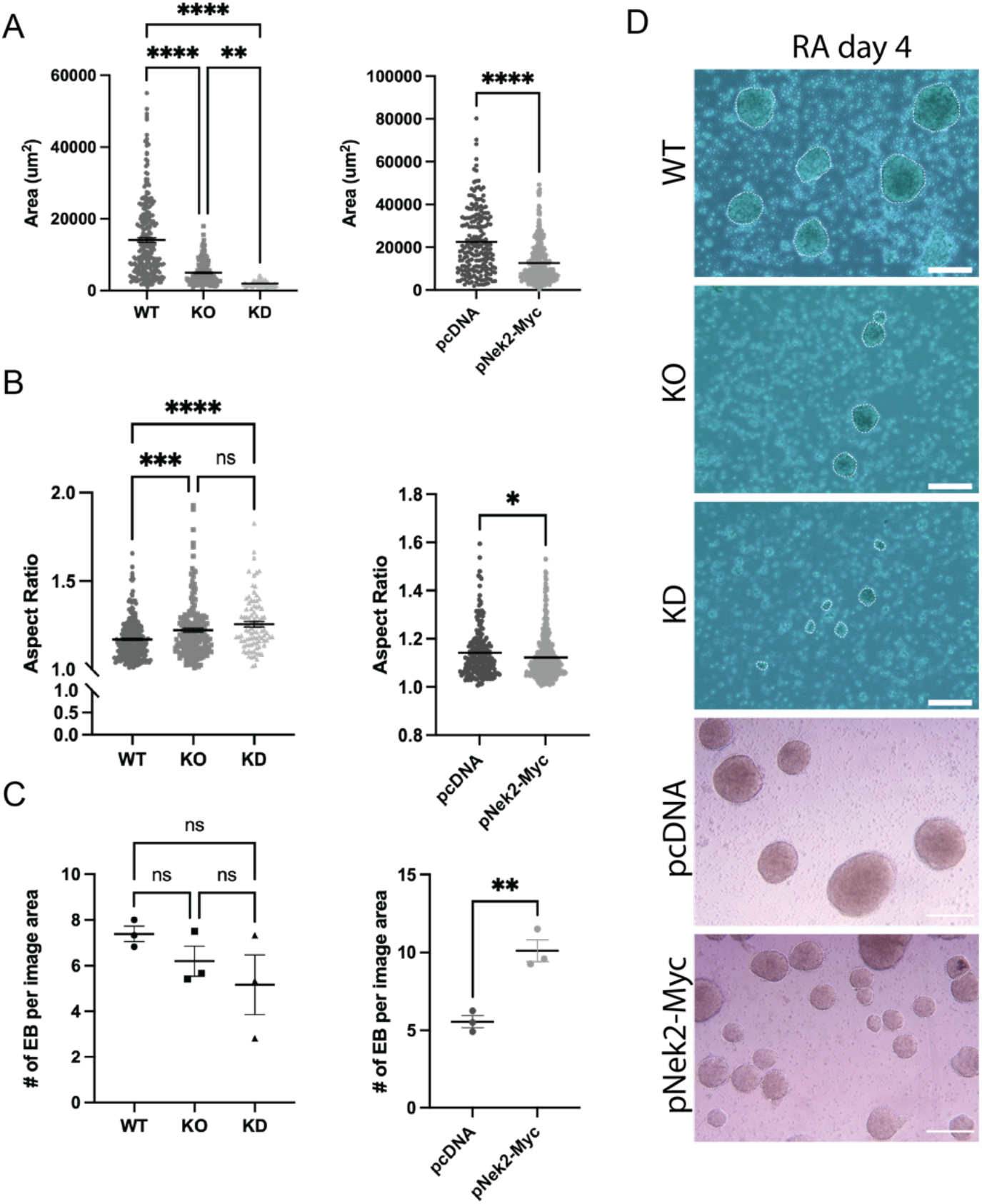
Altering Nek2 levels alters embryoid body development and organization. **(A)** Area, and **(B)** Aspect ratio (major axis/minor axis of EB from images in D formed after 4 days of RA treatment in wildtype (WT), Nek2 knockout (KO) and knockdown (KD) cells, and pcDNA and pNek2-Myc transfected cells. Points represent individual EB. **(C)** Number of EB per image area from images in D. Points represent mean number of EB across all images per replicate. Lines across points represent mean values ±s.e.m. **(D)** Phase contrast images of EB. White dotted lines outline perimeter. Scale bar = 200 um. N=3. P-values were determined by One-way ANOVA with Tukey’s post-hoc analysis. *\*P<0.5, **P<0.01, ***P<0.001, ****P<0.0001*.

### 4.3. Nek2 promotes exit from pluripotency

Since Nek2 has been explored sparingly in the context of pluripotency [47] but as shown can affect cell number, the expression and protein abundance of various factors was examined to determine if there was a relationship. RT-qPCR was used to measure the expression of known pluripotency transcription factors *Oct4* and *Nanog* [48] in undifferentiated WT, KO and KD cells and in those overexpressing *Nek2* compared to a pcDNA control. No significant change in *Oct4* levels were detected in any condition, but there was a significant increase in *Nanog* expression in KO and KD cells (Fig. 4A). Conversely, and relative to the control, *Nek2* overexpressing cells showed a decrease in *Nanog* levels (Fig. 4B) in the undifferentiated state. KO cells were further explored throughout RA-induced differentiation where expression of *Nanog* was elevated in RA-treated cells at days 10, 14 and 17 compared to WT controls (Fig. 4C). Again, *Nek2* overexpressing cells, relative to the control, had reduced *Nanog* expression at days 0, 10 and 14 of RA treatment (Fig. 4D). ESRRB, a marker of pluripotency [48], was also explored in KO and WT cells throughout RA treatment and results showed that it was significantly higher than the WT controls on day 10 following RA treatment, and significantly higher in KO cells relative to WT at day 0 when RA was not present (Fig. 4E). In support, and relative to the pcDNA controls, *Nek2* overexpressing cells lost the ESRRB signal in untreated cells (Fig. 4F). Thus, markers of pluripotency decreased when *Nek2* was overexpressed and increased with Nek2 loss, suggesting it is likely involved in influencing the state of cell pluripotency.

**Figure 4.**
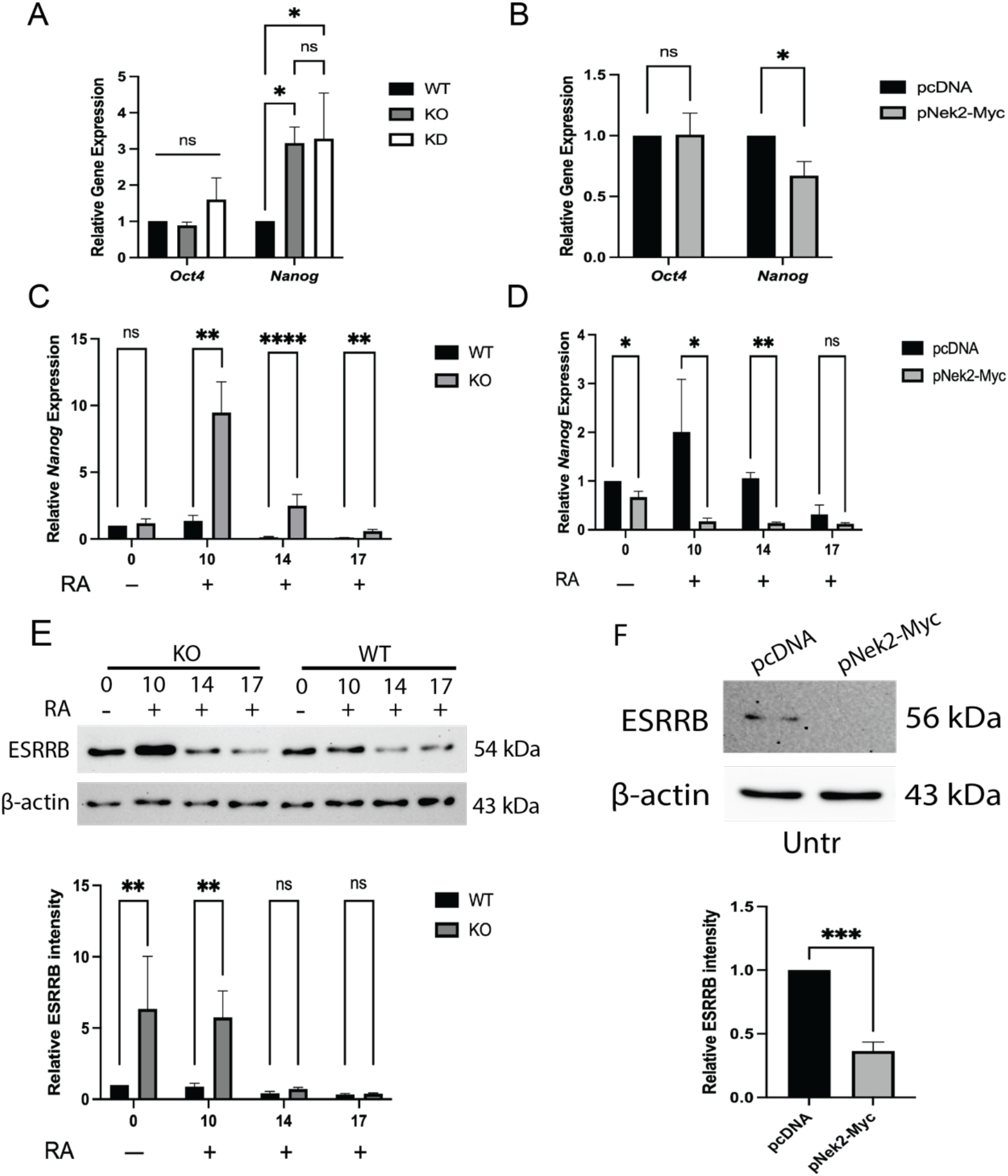
Nek2 promotes exit from pluripotency. RT-qPCR of pluripotency markers *Oct4* and *Nanog* in untreated, undifferentiated **(A)** wildtype (WT), Nek2 knockout (KO) and knockdown (KD) cells, and **(B)** pcDNA and pNek2-Myc transfected cells. P-values were determined by One-way ANOVA with Tukey’s post-hoc analysis. Expression of *Nanog* in **(C)** WT and KO cells and **(D)** pcDNA and pNek2-Myc transfected cells during 0-17 days of RA treatment. Immunoblotting of ESRRB in (E) WT and KO cells during days 0-17 of RA treatment and (F) untreated pcDNA and pNek2-Myc transfected cells. P-values were determined by Two-way ANOVA with Sidak’s post-hoc analysis or by Student’s t-test. N=3. Bars represent mean values ±s.e.m. *\*P<0.05, **P<0.01, ***P<0.001, ****P<0.0001*.

### 4.4. Nek2 is required for neural differentiation

To test the involvement of Nek2 in neural differentiation, antibodies to β-III-tubulin and GFAP were used to detect the presence of neurons and astrocytes, respectively, in WT and Nek2 deficient cells (KO and KD) and in cells overexpressing *Nek2* (pNek2-Myc) (Fig. 5). Compared to WT cells, both markers showed either reduced or absent β-III-tubulin (Fig. 5A and B, Supplemental Figure 4A and B) and GFAP levels (Fig. 5A and C, Supplemental Figure 4A and C) at days 10, 14 and 17 in KO and KD cells. Overexpression of *Nek2* caused the opposite to occur for β-III-tubulin, showing a significant increase at days 10 and 17 (Fig. 5A and B), while GFAP levels were absent in these cells at day 17 (Fig. 5A). Thus, the reduction or absence of Nek2 was sufficient to attenuate both neuronal and astrocyte cell fates. More importantly, overexpressing *Nek2* was sufficient to enhance the neuronal fate, but it also attenuated astrocyte formation. Immunofluorescence was used to corroborate the immunoblot findings at day 17 of RA treatment, and results confirmed no detection of either marker in KO cells (Fig. 5D). To test whether or not Nek2 KO cells remained stem-like in nature, RT-qPCR was used to detect *Nestin*, a neural stem cell marker [49–51]. Results, compared to undifferentiated controls, showed that *Nestin* expression increased in both wildtype and KO cell lines over time (*P<0.0001*), however, there was no significant difference between WT and KO cells at any of the timepoints tested (Fig. 5E). Thus, it appears Nek2 is required for the differentiation of neurons and astrocytes, and its overexpression promotes the formation of neuron but not astrocytes.

**Figure 5.**
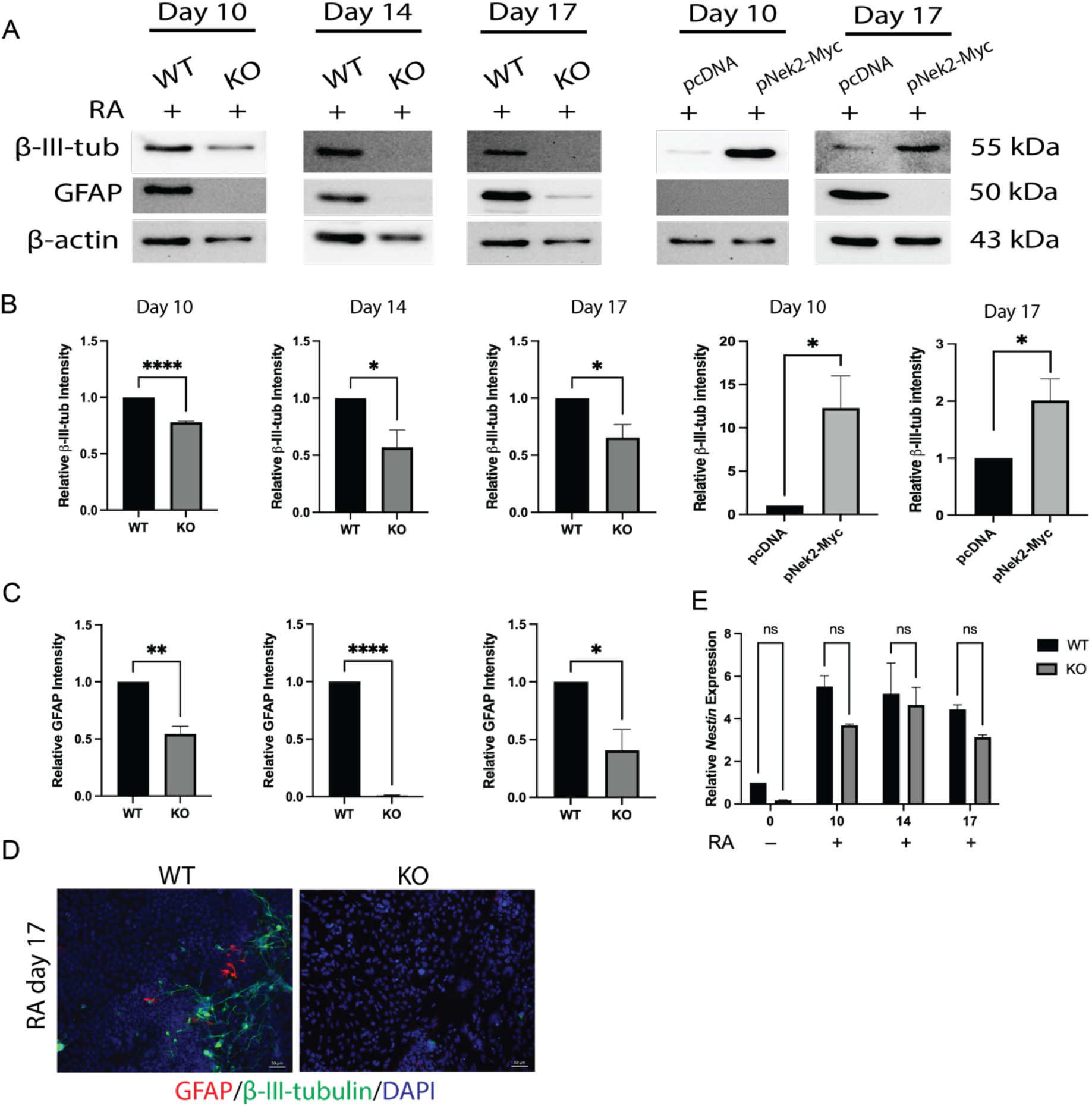
Nek2 is required for neural differentiation. **(A)** Immunoblot of neuron marker, β-III-tubulin, and astrocyte marker, GFAP, in wildtype (WT), Nek2 knockout (KO), pcDNA and pNek2-Myc transfected cells at various times during RA treatment. Densitometry of **(B)**β-III-tubulin and **(C)** GFAP from immunoblots in A). P-values were determined by Student’s t-test. **(D)** Immunofluorescence of GFAP (red), β-III-tubulin (green), and DAPI (blue) staining in WT and KO cells at day 17 of RA treatment. **(E)** RT-qPCR of neural stem cell marker *Nestin* in WT and KO cells during days 0-17 of RA treatment. P-values were determined by Two-way ANOVA with Sidak’s post-hoc analysis. N=3. Bars represent mean values ±s.e.m. *\*P<0.05, **P<0.01, ****P<0.0001*.

### 4.4 Nek2 loss and overexpression alters Hh signalling

Canonical Hh and Wnt signalling both play an integral role in neural differentiation of P19 cells and have been linked to Nek2 [20–23]. Wang et al. [19] and Zhou et al. [20] reported Nek2 serves as a negative regulator of Hh signalling and our previous report has shown that overactivation of Hh signalling through the loss of SUFU inhibited astrocyte differentiation in the P19 model [37] (Spice et al., 2022, under revision). Therefore, Hh signalling was first explored as a potential mechanism for Nek2 in regulating neural differentiation. RT-qPCR was performed in untreated, undifferentiated KO and KD cells, where Hh target gene *Ptch1* was increased compared to WT cells (Supplemental Figure 5A). However, Hh targets *Gli1* and *Ascl1* were only elevated in KD and not KO cells (Supplemental Figure 5A). A Gli-responsive luciferase assay was also performed, which confirmed that untreated KO and KD cells had similar Gli-luciferase activity compared to WT cells treated with 10 nM SAG (Supplemental Figure 5B). Despite the Nek2 KO and *Nek2* overexpression in cells showing opposite levels of pluripotency genes (Fig. 4), there was also an increase in *Ptch1* expression in pNek2-Myc transfected, undifferentiated cells (Supplemental Figure 5C). To get a better understanding how Hh signalling may be affected by altering Nek2 levels, RT-qPCR was also performed on Nek2-deficient and *Nek2* overexpressing cells throughout RA treatment (Fig. 6). Nek2 KO cells showed increased expression of *Gli1* at days 10 and 14 of RA treatment (Fig. 6A), whereas *Nek2* overexpressing cells showed decreased expression at day 10 (Fig. 6B). In Nek2 KO cells, *Ptch1* expression compared to WT controls occurred only at day 0 (Fig. 6C) and at later stages its amount was like that in controls. However, *Nek2* overexpressing cells had reduced expression compared to the controls at day 10 of RA treatment (Fig. 6D). Together, these data suggest that the presence of Nek2 acts to reduce Hh target gene expression at these later stages, however, the link between Nek2 and Hh remains unclear as the expression of target genes was not consistent when Nek2 levels were perturbed (Figure 6).

**Figure 6.**
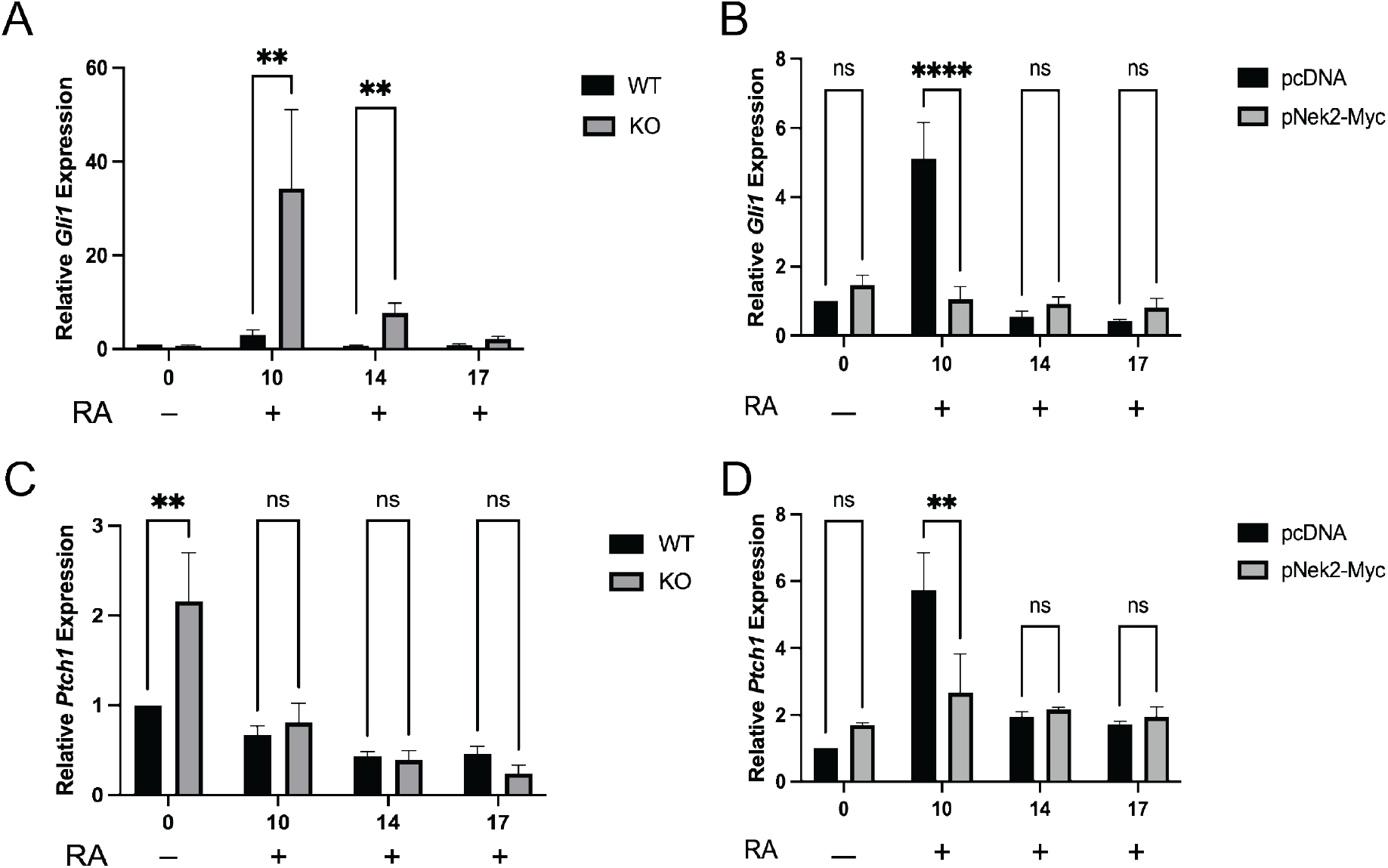
Hh signaling is affected by perturbed Nek2 levels. RT-qPCR of Hh target gene *Gli1* in **(A)** wildtype (WT) and Nek2 knockout (KO) cells, and **(B)** in pcDNA and pNek2-Myc transfected cells during days 0-17 of RA treatment. RT-qPCR of Hh target gene *Ptch1* in **(C)** WT and KO, and **(D)** pcDNA and pNek2-Myc transfected cells during days 0-17 of RA treatment. P-values were determined by Two-way ANOVA with Sidak’s post-hoc analysis. N=3. Bars represent mean values ±s.e.m. *\*\*P<0.01, ****P<0.0001.*

### 4.5. Nek2 loss and overexpression alters Wnt signalling

Since Nek2 is reported as a positive regulator of Wnt signalling through phosphorylation of DVL and β-catenin [20,21], we also explored if the canonical Wnt pathway was linked to Nek2 and neural differentiation. RT-qPCR was performed in untreated, undifferentiated WT, KO and KD cells where the Wnt target gene *Dab2* showed no change in expression. However, other targets including *Dkk1* and *c-Myc* were reduced in KO cells and elevated in KD cells (Supplemental Figure 5D). To better understand these differences, experiments with a β-catenin-responsive luciferase assay was performed in untreated WT, KO and KD cells and in WT cells treated with 10 nM BIO after 24 hours. Results showed KO and KD cells had greater β-catenin-luciferase activity than WT cells, and comparable to when BIO was included as a treatment of WT cells (Supplemental Figure 5E). Significantly, *Nek2* overexpressing cells also showed an increase in the expression of *Dkk1* and *Dab2* compared to that in controls (Supplemental Figure 5F). Furthermore, we employed a similar strategy as that conducted for Hh signalling to investigate the potential link between Wnt and Nek2 at later stages involving RA-induced differentiation. RT-qPCR showed that target gene *Dkk1* was increased at day 10 in KO cells compared to WT controls (Fig. 7A), while *Nek2* overexpressing cells showed consistent increases in *Dkk1* expression at days 0, 14 and 17 of RA treatment; the apparent increase on day 10 was not significant (Fig. 7B). *Dab2* was also investigated, where KO cells showed significantly reduced expression only at day 17 compared to WT cells (Fig. 7C). In contrast, when Nek*2* was overexpressed in cells, *Dab2* was increased at days 14 and 17 of RA treatment (Fig. 7D). Thus, it appeared that the link between Wnt signalling and *Nek2* in regulating neuronal development was clearer than Hh but still not definitive.

**Figure 7.**
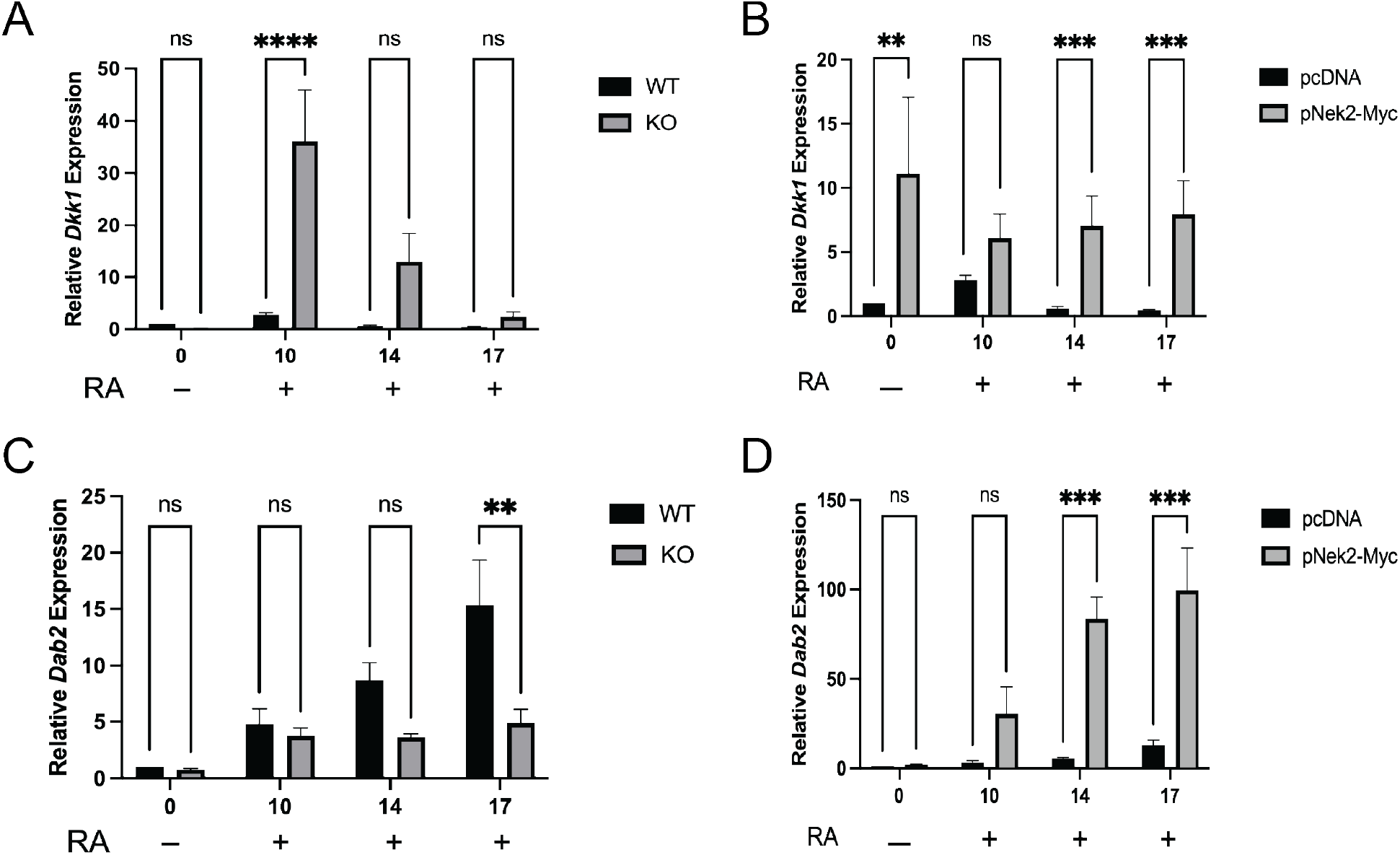
Wnt signaling is affected by perturbed Nek2 levels. RT-qPCR of Wnt target gene *Dkk1* in **(A)** wildtype (WT) and Nek2 knockout (KO) cells, and **(B)** in pcDNA and pNek2-Myc transfected cells during days 0-17 of RA treatment. RT-qPCR of Hh target gene *Dab2* in **(C)** WT and KO, and **(D)** pcDNA and pNek2-Myc transfected cells during days 0-17 of RA treatment. P-values were determined by Twoway ANOVA with Sidak’s post-hoc analysis. N=3. Bars represent mean values ±s.e.m. *\*\*P<0.01, ***P<0.001, ****P<0.0001*.

Since *Wnt1* overexpression in P19 cells promotes neuron differentiation while inhibiting astrocyte differentiation [3] we employed a chemical strategy to further test this relationship between Nek2 and Wnt in neurogenesis. P19 cells were treated with BIO [52], an inhibitor of GSK3 that activates the Wnt pathway, and XAV [53], a tankyrase inhibitor which serves as a Wnt pathway antagonist. Co-treating cells with RA and BIO did not alter β-III-tubulin levels relative to RA alone (Supplemental Figure 6A and B), or when treated first with BIO before the addition of RA four days later. RA was the key inducer as BIO alone showed little to no β-III tubulin levels. The levels of GFAP, however, was greatly affected at later stages when the canonical Wnt pathway was activated by BIO in RA-treated cells (Supplemental Figure 6A and C). XAV treatment, expected to produce opposite results, showed cells co-treated with it and RA inhibited both β-III-tubulin and GFAP detection (Supplemental Figure 6D-F). Together, these chemical and genetic results reveal a link between Nek2 and Wnt signalling, specifically on astrocyte differentiation, but the mechanism is still unclear on how the loss of Nek2 attenuates neuronal differentiation.

### 4.6. Proteomic analysis identifies several differentially abundant proteins in Nek2 deficient cells

Although canonical Hh and Wnt signalling is required in P19 cells for proper neural differentiation [3,37,40,54], and there are relationships between these pathways and Nek2, our data strongly suggests that the loss of Nek2 influences neural cell lineages linked to pluripotency and cell proliferation. The data supports this early role, rather than being directly linked to regulating the Wnt and Hh signalling pathways. To investigate this further a proteomic approach comparing global protein abundance patterns was chosen to provide more information of the pathways involving Nek2 in P19 cells. Proteins were analyzed from cells collected at days 0, 1, 4 and 10 of RA induced differentiation of WT and KO cells (Supplemental Figure 7). Differential enrichment analysis identified 1780 differentially regulated proteins across all time points (Supplemental Figure 7C-F); however, only 3 proteins were upregulated in KO cells (Fig. 8A): Cth, Slc25a13 and Pla2g15. In exploring pathways enriched specifically at day 10 of RA treatment, KO cell pathway enrichment included proteins involved in fatty acid metabolism, the G2 DNA damage checkpoint and K63 linked deubiquitination (Fig. 8C). Sixteen proteins were found to be downregulated in KO cells (Fig. 8B), including Ndufb8, Ndufv1, Ndufa10, Ndufc2, Mtnd4, Ptgr1, Mtnd1, Ndufs1, Mtco2, Ndufa4, Ndufa12, Ndufa5, Ndufb1, Cox6b1, Mtnd5 and Gstp1, and pathway enrichment in WT cells included proteins involved in negative regulation of cell projection organization, negative regulation of neuron projection development and mitochondria localization (Fig. 8D). Other pathways enriched in the WT population include Myc targets, epithelial to mesenchymal transition proteins and those related to oxidative phosphorylation (Supplemental Figure 7G).

**Figure 8.**
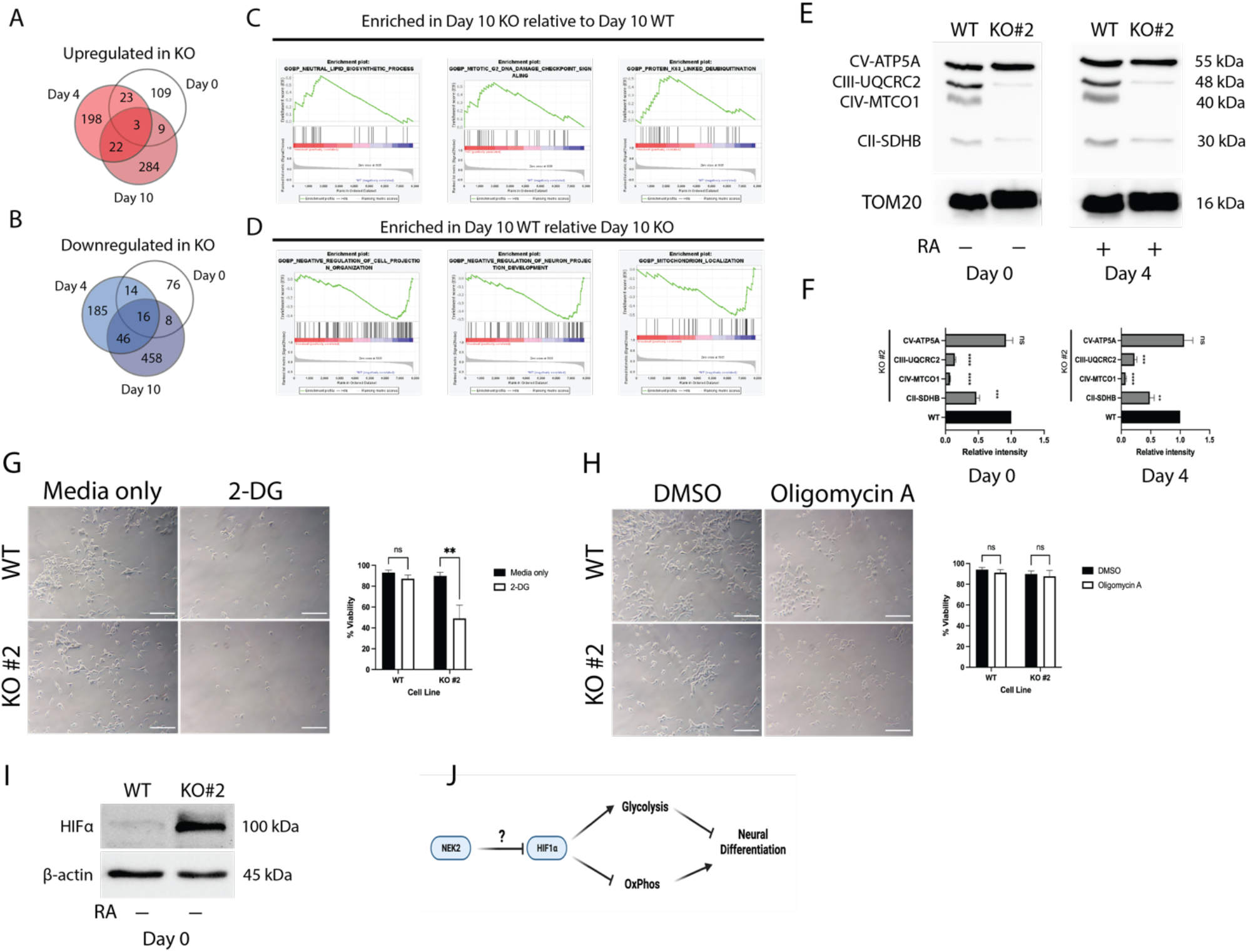
Loss of Nek2 inhibits oxidative metabolism. Proteins identified through LC-MS that are **(A)** upregulated and **(B)** downregulated in KO cells compared to WT cells at days 0, 4 and 10 of RA treatment. GSEA enrichment analysis of LC-MS results identifying pathways that are **(C)** enriched in KO cells compared to WT cells and **(D)** in WT cells compared to KO cells at day 10 of RA treatment. **(E)** Immunoblot of electron transport chain components, SDHB, MTCO1, UQCRC2 and ATP5A corresponding to subunits in complexes II-V, respectively, in WT and KO cells at days 0 and 4 of RA treatment. **(F)** Densitometry analysis of immunoblots in E. P-values were determined by One-way ANOVA with Tukey’s post-hoc analysis. Phase contrast images and viability of cells determined through trypan blue exclusion in cells treated with **(G)** media only or 50 mM 2-DG, and **(H)** DMSO or 2.5 μM oligomycin A for 24 hours. Scale bar = 200 μm. P-values were determined by Two-way ANOVA with Sidak’s post-hoc analysis. (I) Immunoblot of HIF1α in untreated WT and KO cells. (J) Proposed mechanism of Nek2 action. Image created using BioRender. Bars represent mean values ±s.e.m. *\*\*P<0.01, ***P<0.001, ****P<0.0001*.

### 4.7. Loss of Nek2 promotes glycolytic metabolism

Since most proteins negatively impacted in Nek2 KO cells involved complexes of the mitochondrial electron transport chain (ETC) and that WT cells compared to KO cells at day 10 had enriched proteins related to oxidative phosphorylation (OxPhos; Supplemental Figure 7), we wanted to test what metabolic pathway was being utilized in Nek2 KO cells. Immunoblotting was used and like the LC-MS results, results showed KO cells at days 0 and 4 of RA treatment had reduced or absent ETC complexes II, III and IV (Fig. 8E and F). Furthermore, these cells were treated with either 50 mM 2-deoxy-D-glucose (2-DG) to inhibit glycolysis, or 2.5 μM oligomycin A to inhibit OxPhos to determine if, compared to WT cells, they were reliant on one or the other metabolic pathways. Results showed WT cell viability was unaffected by either treatment (Fig. 8G and H, respectively) suggesting these cells likely have a bivalent metabolism in the undifferentiated state. However, 2-DG significantly reduced KO cell viability (Fig. 8H), implying these cells rely on glycolysis. As glycolytic genes are known to be upregulated in response to hypoxia-inducible factor 1α (HIF1α) stabilization [55], we explored HIF1α levels in untreated, undifferentiated WT and KO cells (Fig. 8I). Interestingly, under normoxic conditions, undifferentiated WT cells do not show HIF1α stabilization like that seen in KO cells (Fig. 8I). These results show that Nek2 is essential for maintaining ETC components and that the loss of Nek2 shifts P19 cells towards glycolysis and HIF1α stabilization (Fig. 8J).

## 5. Discussion

Nek2 has many roles within the cell and is involved in regulating centrosome dynamics, signalling pathways and metabolism, and although its importance is clear, few studies have explored this protein within normal development [24–26]. We have shown that Nek2 is essential in the differentiation of both neurons and astrocytes in the P19 embryonal carcinoma model (Fig. 5) and previous reports have placed Nek2 as a regulator of the Hh and canonical Wnt pathways [20–23,27], which are required in this process [3,5,37–40] (Supplemental Figure 6). Towards that end, we sought to explore neural differentiation using the P19 model through the context of Wnt and Hh signalling, and although Nek2 appears to affect these pathways, the data did not indicate that their deregulation was the causative factor affecting neural differentiation. We did find that a reduced proliferative capacity of cells lacking Nek2 (Fig. 2 and 3) and a metabolic shift away from OxPhos towards exclusively glycolytic metabolism (Fig. 8) as the likely cause of this reduced differentiation potential when Nek2 is absent.

One of the main factors seeming to affect the differentiation of neurons and astrocytes was the reduced proliferative capacity of Nek2 KO cells. Without Nek2, undifferentiated cells had reduced numbers (Fig. 2) as did cells in EBs, where these EBs had reduced size and organization (Fig. 3) without altered cell viability or apoptosis (Supplemental Figure 3). Interestingly, the overexpression of *Nek2* resulted in the opposite phenotype, with increased proliferation in the undifferentiated state (Fig. 2) and reduced EB size, but increased EB number (Fig. 3). The loss of Nek2 was also reported to cause reduced proliferation in glioblastoma and medulloblastoma cell lines, however, this reduction was accompanied by increased apoptosis [29,30]. Another report also observed the reduced ability of Nek2-depleted cells to form spheroids (similar to EBs) in a hepatocellular carcinoma cell line [27]. This study by Lin et al. [27] also showed Wnt3a treatment resulted in an increase in spheroid number in *Nek2* expressing hepatocellular carcinoma cells, which parallels our results that showed when Nek2 was overexpressed it caused canonical Wnt signalling to be activated (Fig. 7) and an increased number of EB (Fig. 3). Despite the well-known phenomenon that pluripotent cells are highly proliferative [56], we were surprised that unlike hepatocellular carcinoma cells depleted of Nek2 [27], our KO cells showed elevated pluripotency factors, *Nanog* and ESRRB (Fig. 4), and reduced proliferation rates (Fig. 2 and 3). These pluripotency factors were also reduced or absent when *Nek2* was overexpressed in P19 cells (Fig. 4), and this was accompanied by an increase in proliferation (Fig. 2). Despite the lack of information reported for Nek2, these differences in pluripotency and proliferation were noted but their significance remains unknown. A previous report in hESCs identified Nek2 activity in the undifferentiated state and during early induction of differentiation [47], which highlights that Nek2 is likely playing a role, however the specifics of its role in pluripotency are unclear. Our results, with this previous report [47], note Nek2 is likely functioning earlier during the “decision-making” process involving pluripotency (Fig. 4) and/or cell fate determination that favours one lineage over the other (Fig. 5). Although we predicted the role of Nek2 lay within regulation of Wnt or Hh, our results point to the regulation of pluripotency and the unlikelihood that Nek2 in P19 cells is directly linked to Hh (Fig. 6) and/or canonical Wnt signalling (Fig. 7) in differentiation.

In addition to changes in pluripotency factors a proteomics approach revealed several proteins with differential enrichment in WT and KO cells and highlighted a reduction in mitochondrial ETC components in Nek2 KO cells (Fig. 8). Further experiments, confirming these LC-MS results, showed that losing Nek2 altered cellular metabolism, with KO cells becoming exclusively glycolytic (Fig 8). Our results are intriguing in the light of the study by Zhou et al. [57] who found in certain types of B-cell lymphomas that Nek2 promotes proliferation but also glycolysis. We explored HIF1α levels and results showed P19 cells lacking Nek2 showed increased HIF1α stabilization (Fig. 8I) and a concomitant reliance on glycolytic metabolism (Fig. 8G). The connection between metabolic changes and neural differentiation has been reported in human induced pluripotent stem cell (hiPSC) derived neural precursors (NPCs) where there is a switch from aerobic glycolysis to OxPhos in neurons involving pyruvate kinase M1/2 (PKM) isoforms [58]. Given that Nek2 is also involved in PKM splicing [57,59] and its loss causes P19 cells to become glycolytic (Fig. 8) and thus blocking neural differentiation (Fig. 5), it seems reasonable to propose its contribution in this process. HIF1α has also been explored within neural differentiation where its overexpression in hiPSCs enhanced pluripotency and reduced neural differentiation [60], which was also observed in mESCs [61]. Previous work with P19 cells have shown that there is a shift from glycolytic metabolism to OxPhos with RA induction and this promotes neuron differentiation [62,63]. Interestingly, and in contrast to our proposed mechanism of cells grown under normoxic conditions (Fig. 8J), a previous report in P19 cells showed an increase in neuron differentiation under hypoxic conditions where HIF1α was stabilized [64]. Given our work, however, we have shown in P19 cells that altered metabolism and neural differentiation is the result of cells lacking Nek2, thus demonstrating an essential role for Nek2 in regulating this metabolic transition from glycolysis to OxPhos.

Evidence would suggest the mechanism of Nek2 regulation likely involves HIF1α, however the direct link between Nek2 and it remains to be established (Fig. 8J). These connections, in addition to the post-translational modifications Nek2 likely performs to promote or inhibit neural differentiation must be further examined and are currently under investigation.

## 6. Acknowledgements

The authors would like to thank past and current members of the Kelly Lab for their support.

## 7. Author contributions

Conceptualization, D.M.S., G.M.K,; Methodology, D.M.S, T.C., G.A.L, G.M.K.; Software, T.C.; Formal Analysis, D.M.S., T.C.; Investigation, D.M.S., T.C.; Resources, G.A.L., G.M.K.; Data Curation, T.C., G.A.L.; Writing – Original Draft, D.M.S., T.C.; Writing – Review & Editing, D.M.S., G.M.K.; Visualization, D.M.S., T.C.; Supervision, G.A.L., G.M.K.; Project Administration, D.M.S., G.M.K.; Funding Acquisition, G.M.K.

