## Supplemental Table and Figures for "Never in Mitosis Kinase 2 regulation of metabolism is required for neural differentiation"

**Supplemental Table 1. CRISPR-Cas9 gRNAs, primers for gRNA amplification and PCR around the cut site.**

| CRISPR-Cas9 | | |
|---|---|---|
| *Nek2* KO gRNA | CATCGTAATGGAATACTGTGAGG | |
| *Nek2* KD gRNA | CAAGCCATCAGAGTAGCGGTAGG | |
| | Forward Primer (5′ → 3′) | Reverse Primer (5′ → 3′) |
| *Nek2* KO gRNA | CACCCATCGTAATGGAATACTGTGAGG | AAACCCTCACAGTATTCCATTACGATG |
| *Nek2* KO PCR | TTTGTCTCACACCTTGGAAATG | TTATTAGATACGAACCGTGGGG |
| *Nek2* KD gRNA | CACCCAAGCCATCAGAGTAGCGGTAGG | AAACCCTACCGCTACTCTGATGGCTTG |
| *Nek2* KD PCR | CTGCAGAATGTCCAGAGCGT | CTGTAGCCAGGGAAACCGAT |
| *U6* | GGGCAGGAAGAGGGCCTAT | |

**Supplemental Table 2. Primers used for RT-qPCR**

| **qRT-PCR** | | |
|---|---|---|
| **Gene** | Forward Primer (5′ → 3′) | Reverse Primer (5′ → 3′) |
| *L14* | GGGTGGCCTACATTTCCTTCG | GAGTACAGGGTCCATCCACTAAA |
| *Gli1* | GGAAGTCCTATTCACGCCTTGA | CAACCTTCTTGCTCACACATGTAAG |
| *Ptch1* | AAAGAACTGCGGCAAGTTTTTG | CTTCTCCTATCTTCTGACGGGT |
| *Oct4* | CCCAATGCCGTGAAGTTGGA | GCTTTCATGTCCTGGGACTCCT |
| *Nanog* | TCTTCCTGGTCCCCACAGTTT | GCAAGAATAGTTCTCGGGATGAA |
| *Dkk1* | TGAAGATGAGGAGTGCGGCTC | GGCTGTGGTCAGAGGGCATG |
| *Dab2* | GGAGCATGTAGACCATGATG | AAAGGATTTCCGAAAGGGCT |

**Supplementary Table 3. Q Exactive Settings for Data Acquisition**

| QE Plus Setting | Value |
|---|---|
| Scan Range | 400-1500 m/z |
| MS1 AGC Target | 3E6 |
| MS1 Resolution | 70K |
| MS2 Resolution | 17.5K |
| MS2 AGC Target | 2E5 |
| Maximum IT | 64ms |
| Loop Count | 12 |
| Top N | 12 |
| Isolation Window | 1.2 m/z |
| Isolation Offset | 0.5 m/z |
| NCE | 25 |
| Dynamix Exclusion | 30 |
| Charge Exclusion | 1,7,8,>8 |
| Fixed First Mass | 100 m/z |

Supplemental Figure 1

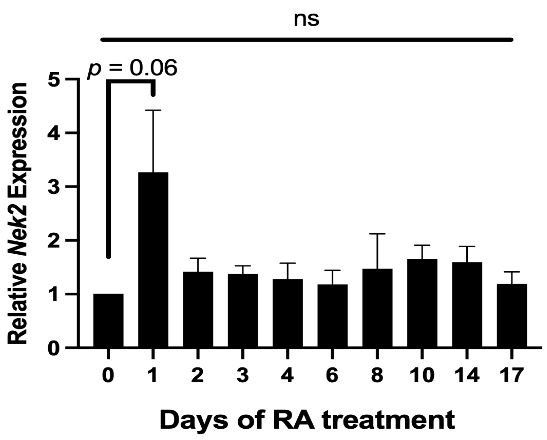


**Supplemental Figure 1. *Nek2* gene expression does not change throughout RA-induced neural differentiation**. RT-qPCR of *Nek2* during days 0-17 of RA treatment. N=3. Bars represent mean values +/- s.e.m. Significance was determined by One-way ANOVA with Tukey's post-hoc analysis.

### Supplemental Figure 2

A

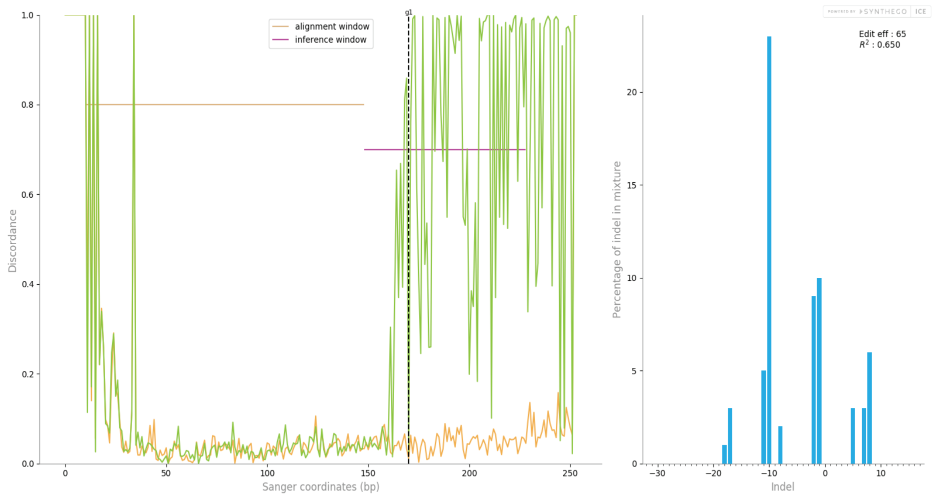


B

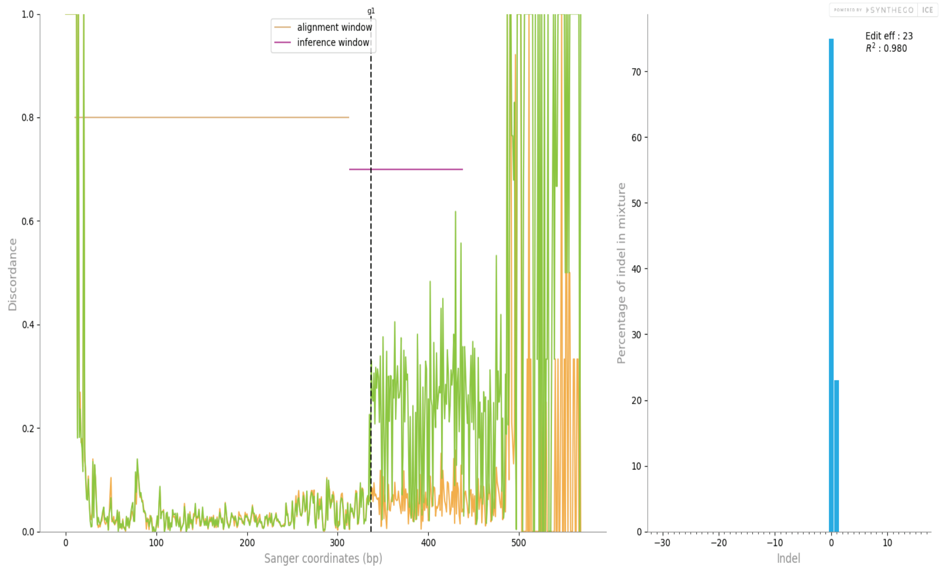


**Supplemental Figure 2. Allele summary of NEK2 deficient cells.** Synthego ICE analysis showing alleles in the population of (A) Nek2 knockout (KO) and (B) Nek2 knockdown (KD) cells. Orange lines show WT sequence similarity to projected region, green lines show mutant sequence similarity. Dotted black line higlights the predicted cut site. Blue bars represent percentage of identified pool with the noted in/del.

Supplemental Figure 3

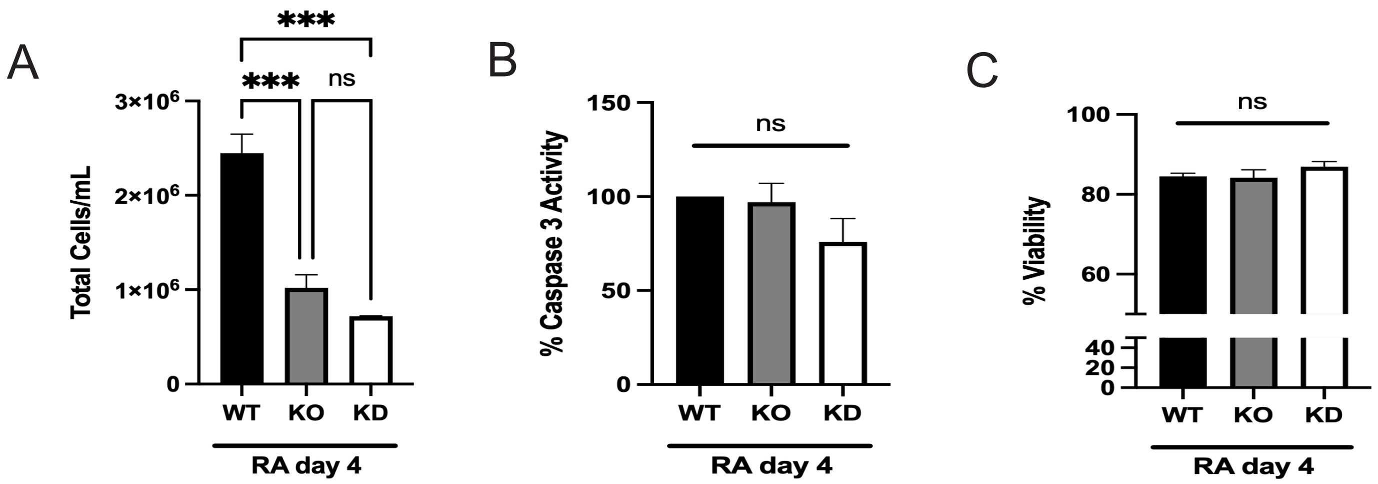


**Supplemental Figure 3. Nek2 deficiency inhibits proliferation without affecting apoptosis. (A)** Total number of cells (F) Caspase 3 activity measured by cytoplasmic concentration of pNA after EBs being resuspended and (C) trypan blue exclusion test measuring cell viability in EB in wildtype (WT), Nek2 knockout (KO) and knockdown (KD) cells. N=3. Bars represent mean values +/- s.e.m. P-values were determined by One-way ANOVA with Tukey's post-hoc analysis. ****P<0.001*, *****P<0.0001*.

### Supplemental Figure 4

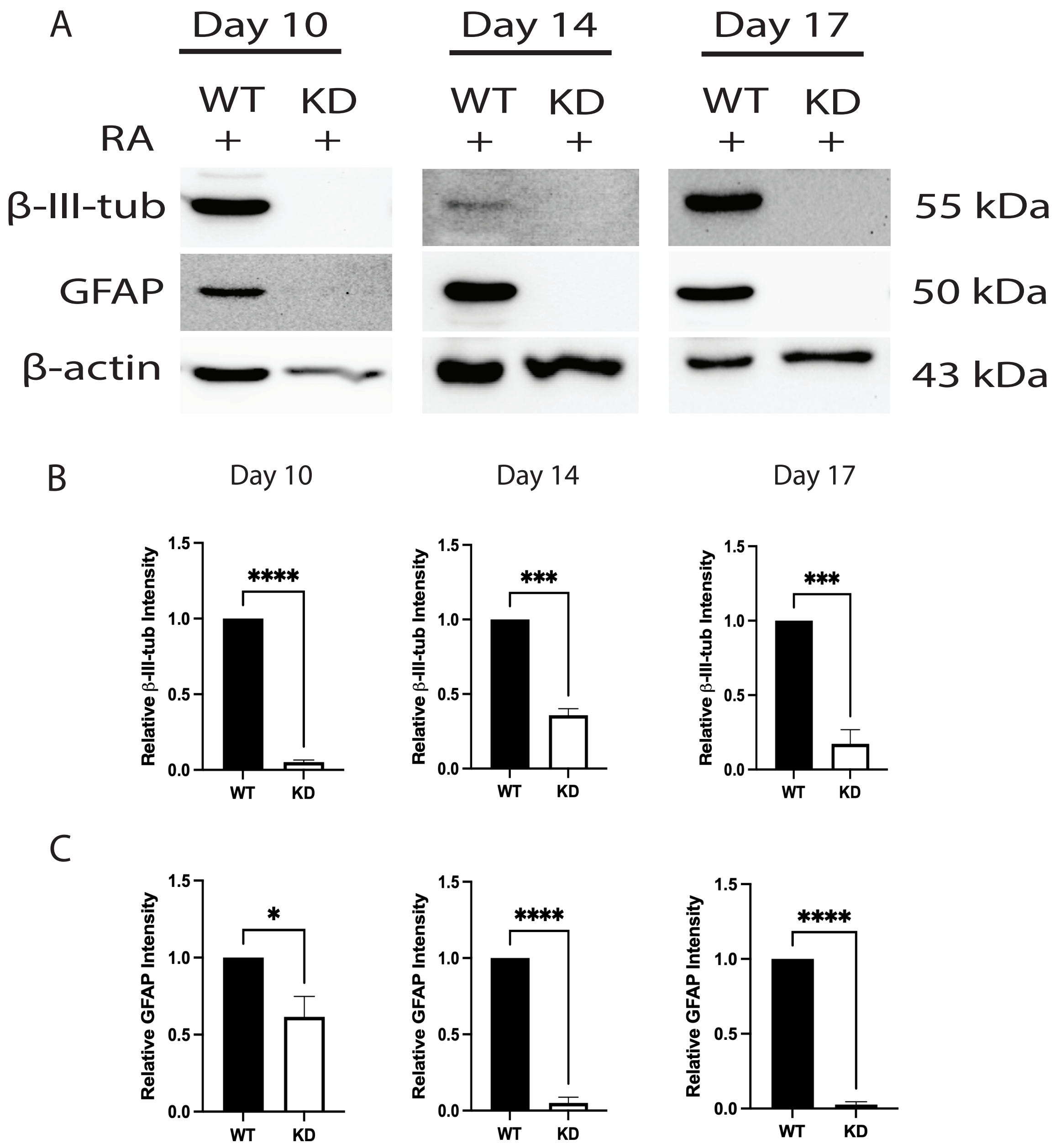


**Supplementary Figure 4. NEK2 knockdown blocks differentiation of neurons and astrocytes. A)** Immunoblot of neuron marker β-III-tubulin, astrocyte marker GFAP and β-actin in wildtype (WT) and *Nek2* knockdown (KD) cells after differentiation in the presence of 0.5 μM retinoic acid (RA) for 10, 14 and 17 days. Densitometry of immunoblot in A of **B)** β-III-tubulin and **C)** GFAP at the same time points. Bars represent mean values +/- s.e.m. N=3. P-values were determined by Student's t-test. **P<0.05, **P<0.01, ***P<0.001, ****P>0.0001*.

### Supplemental Figure 5

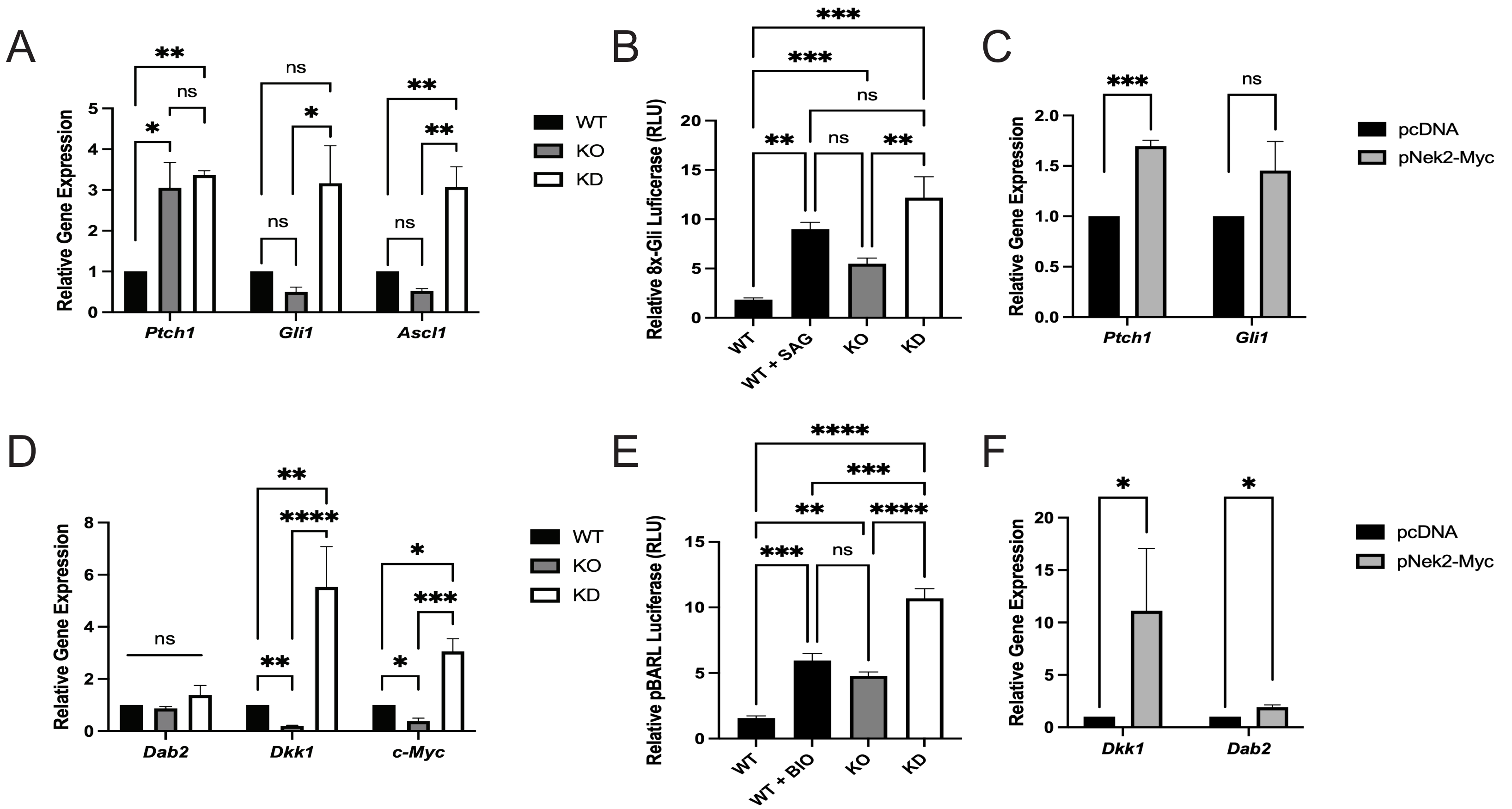


**Supplemental Figure 5. Hh and Wnt signaling are perturbed in Nek2 deficient and overexpressing cells. (A)** RT-qPCR of Hh target genes *Ptch1*, *Gli1* and *Ascl1* in wildtype (WT), knockout (KO) and knockdown (KD) cells in undifferentiated cells. **(B)** Gli-responsive luciferase assay in untreated WT, KO and KD cells and WT cells with 10 nM SAG treatment after 24 hours. **(C)** RT-qPCR of Hh target gene *Ptch1* and *Gli1* in pcDNA and pNek2-Myc transfected cells. **(D)** RT-qPCR of Wnt target genes *Dab2*, *Dkk1* and *c-Myc* in WT, KO and KD undifferentiated cells. **(E)** β-catenin responsive luciferase assay in untreated WT, KO and KD cells and WT cells treated with 10 nM BIO after 24 hours. **(F)** RT-qPCR of Wnt target genes *Dkk1* and *Dab2* in pcDNA and pNek2-Myc transfected cells. P-values were determined by One-way ANOVA with Tukey's post-hoc analysis. N=3. Bars represent mean values +/- s.e.m. **P<0.05*, ***P<0.01*, ****P<0.001*, *****P<0.0001*.

### Supplemental Figure 6

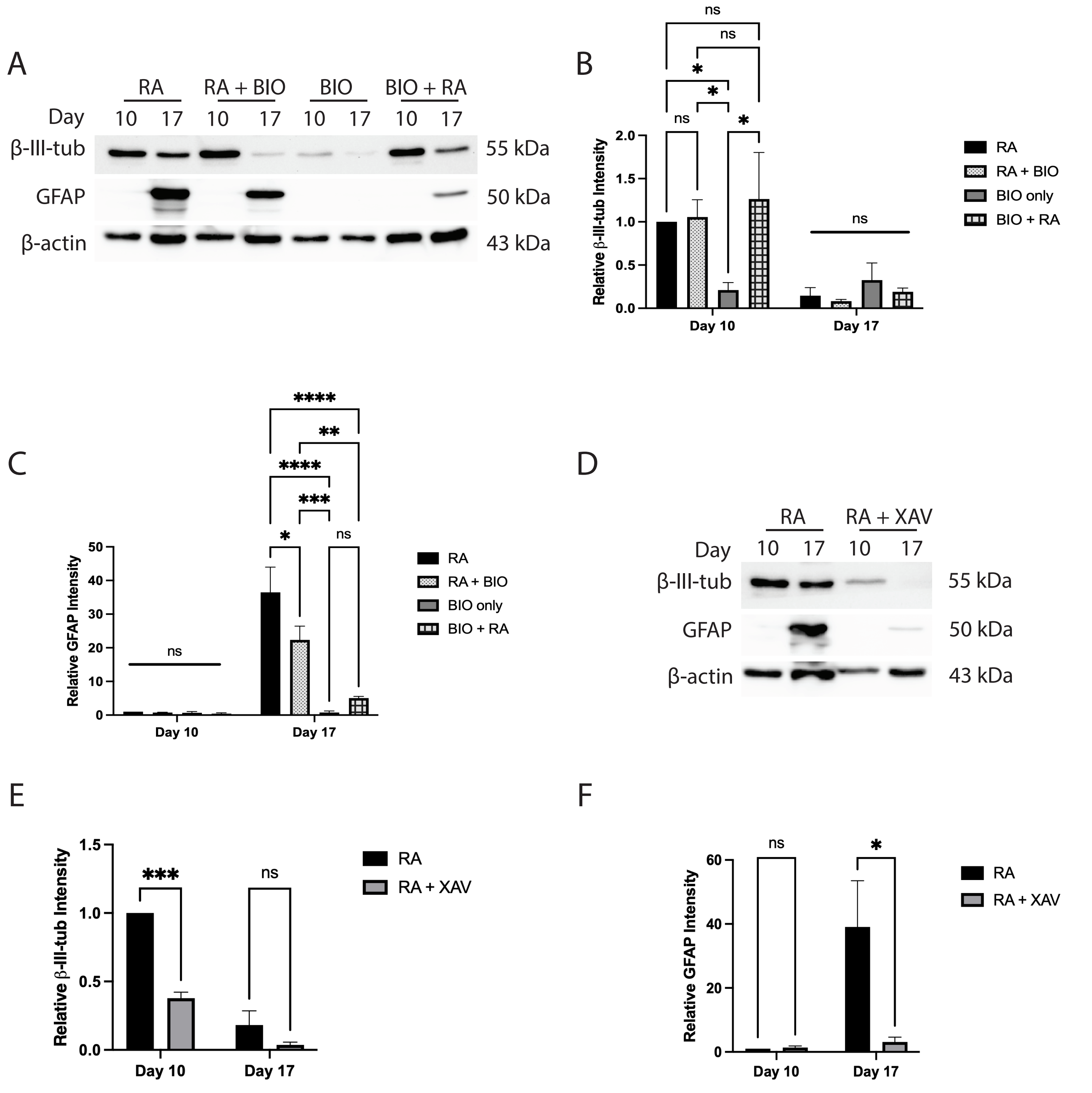


**Supplemental Figure 6. Wnt is required for neural differentiation but is only sufficient to induce neurons. (A)** Immunoblot of ß-III-tubulin and GFAP in wildtype cells at days 10 and 17 after after treatment with 0.5 uM on days 0 and 4 (RA), 0.5 uM RA with 10 nM BIO on day 0 plus RA on day 4 (RA+BIO), 10 nM BIO treatment on day 0 (BIO) or 10 nM BIO treatment on day 0 plus RA on day 4 (BIO+RA). Densitometry analysis of the immunoblot in A for **(B)** ß-III-tubulin and **(C)** GFAP. **(D)** Immunoblot of ß-III-tubulin and GFAP in wildtype cells at days 10 and 17 after treatment with 0.5 uM RA on days 0 and 4 (RA) or 0.5 uM RA with 10 uM XAV on day 0 plus RA on day 4. Densitometry analysis of the immunoblot in D of **(E)** ß-III-tubulin and **(F)** GFAP. N=3. Bars represent mean values +/- s.e.m. P-values were determined by Two-way ANOVA with Sidak's post-hoc analysis. **P<0.05*,***P<0.01*, ****P<0.001*, *****P<0.0001*.

### Supplemental Figure 7

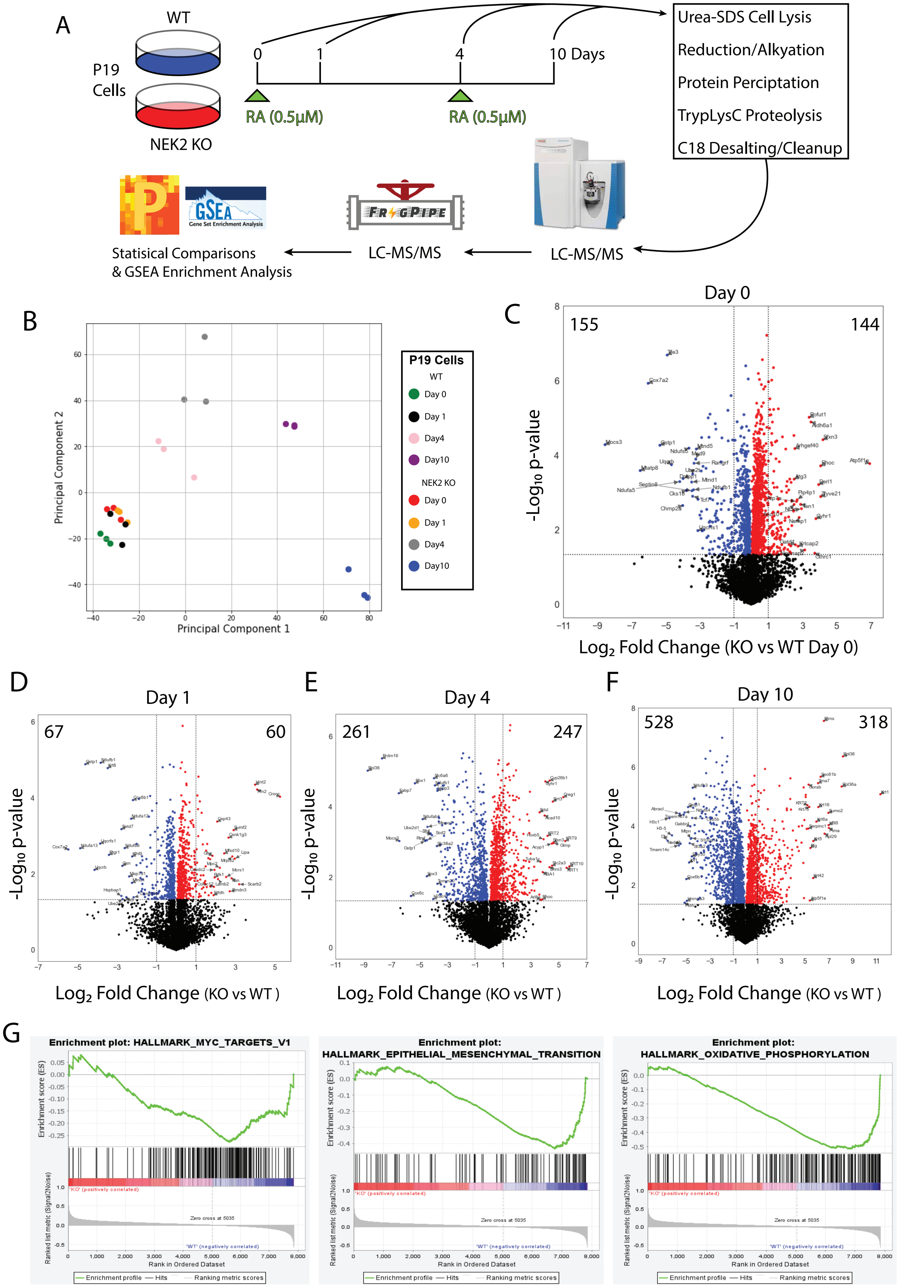


**Supplemental Figure 7. LC-MS workflow and results of WT and KO cells. (A)** Workflow of LC-MS. **(B)** Principle component analysis of samples used for LC-MS showing clustering of biological replicates. Proteins identfied as up or down regulated in KO cells compared to WT cells at days **(C)** 0, **(D)** 1, **(E)** 4 and **(F)** 10 of RA treatment. **(G)** GSEA Pathway Enrichment Analysis of proteins enriched in WT cells compared to KO cells.
